# DISEASES: Text mining and data integration of disease–gene associations

**DOI:** 10.1101/008425

**Authors:** Sune Pletscher-Frankild, Albert Pallejà, Kalliopi Tsafou, Janos X. Binder, Lars Juhl Jensen

**Affiliations:** Novo Nordisk Foundation Center for Protein Research, Faculty of Health and Medical Sciences, University of Copenhagen; Novo Nordisk Foundation Center for Basic Metabolic Research, Faculty of Health and Medical Sciences, University of Copenhagen, Copenhagen, Denmark; Structural and Computational Biology Unit, European Molecular Biology Laboratory (EMBL), Heidelberg, Germany; Bioinformatics Core Facility, Luxembourg Centre for Systems Biomedicine (LCSB), University of Luxembourg, Luxembourg

## Abstract

Text mining is a flexible technology that can be applied to numerous different tasks in biology and medicine. We present a system for extracting disease–gene associations from biomedical abstracts. The system consists of a highly efficient dictionary-based tagger for named entity recognition of human genes and diseases, which we combine with a scoring scheme that takes into account co-occurrences both within and between sentences. We show that this approach is able to extract half of all manually curated associations with a false positive rate of only 0.16%. Nonetheless, text mining should not stand alone, but be combined with other types of evidence. For this reason, we have developed the DISEASES resource, which integrates the results from text mining with manually curated disease–gene associations, cancer mutation data, and genome-wide association studies from existing databases. The DISEASES resource is accessible through a user-friendly web interface at http://diseases.jensenlab.org/, where the text-mining software and all associations are also freely available for download.

## Introduction

### Named entity recognition (NER)

Recognizing named entities and concepts, such as genes and diseases, in text is the basis for most biomedical applications of text mining [1]. NER is sometimes divided into two subtasks, namely recognition and normalization (also known as identification or grounding), the former being to recognize the words of interest and the latter being to map them to the correct identifiers in databases or ontologies. However, as recognition without normalization has very limited practical use, the normalization step is now often implicitly considered part of the NER task.

The main challenges in NER are the poor standardization of names and the fact that a name of, for example, a gene or disease may have other meanings [2]. To recognize names in text, many systems thus make use of rules that look at features of names themselves, such as capitalization and word endings, as well as contextual information from nearby words. In early methods the rules were hand crafted [3], whereas newer methods make use of machine learning [4,5], relying on the availability of manually annotated text corpora.

Dictionary-based methods instead rely—as the name suggests—on matching a dictionary of names against text. For this purpose the quality of the dictionary is obviously very important; the best-performing methods for NER according to blind assessments rely on carefully curated dictionaries to eliminate synonyms that give rise to many false positives [6,7]. Moreover, dictionary-based methods have the crucial advantage of being able to normalize names. Whether or not one makes use of machine learning, a high-quality, comprehensive dictionary of gene and disease names is thus a prerequisite for mining disease–gene associations from the biomedical literature.

### Controlled vocabularies of diseases

It is fairly straightforward to find a good starting point for a dictionary of human gene names due to efforts such as the Human Genome Organization (HUGO) Gene Nomenclature Committee (HGNC) [8] and UniProt Knowledgebase (UniProtKB) [9]. It is less obvious to find a good dictionary of disease names, as there are several competing classifications and ontologies, which are designed for different purposes, mutually inconsistent, and thus poorly integrated with each other.

In a clinical setting, various versions of the International Classification of Diseases (ICD; http://www.who.int/classifications/icd/) are almost ubiquitously used for coding diagnoses in electronic health records (EHRs) and derived health registries [10]. European countries, Canada, and Australia use revision 10 (ICD-10), whereas the United States still use revision 9 (ICD-9). ICD-10 is not just an update to ICD-9; it is a restructured diagnosis classification, and no official mapping exists between the two revisions. Because ICD is designed for clinical coding and billing purposes, its structure and disease names are poorly suited for biomedical literature mining. It is, however, useful for text mining of clinical narrative in EHRs, especially because it has been translated to many languages [11].

A newer alternative is the Systematized Nomenclature of Medicine – Clinical Terms (SNOMED CT; http://www.ihtsdo.org/snomed-ct/). It cross maps to several revisions of ICD and has a considerably broader scope than just diseases. SNOMED-CT is one of many terminologies combined in the even broader Unified Medical Language System (UMLS) Metathesaurus; another is Medical Subject Headings (MeSH; http://www.ncbi.nlm.nih.gov/mesh/). Dictionaries based on subsets of UMLS have been used for recognition of disease names with varying success in text-mining tools, such as MetaMap [20442139], Medical Language Extraction and Encoding (MedLEE) [12], and the Mayo clinical Text Analysis and Knowledge Extraction System (cTAKES) [13]. However, because UMLS contains many distinct concepts that are very close in meaning even human annotation of UMLS concepts in text is problematic [14]. Licenses for SNOMED-CT and other terminologies in UMLS further restrict their use in resources intended for redistribution.

In contrast to these, the Disease Ontology [15] is part of the Open Biomedical Ontologies (OBO) Foundry initiative [16]. It cross maps to UMLS and has extensive annotation of synonyms. Consequently, Disease Ontology works well for recognition of diseases in Gene Reference Into Function (GeneRIF; http://www.ncbi.nlm.nih.gov/gene/about-generif) entries [17].

### Information extraction (IE)

Having addressed the NER task using appropriate dictionaries of gene and disease names, the next task is to extract information on associations between genes and diseases. There are two fundamentally different approaches to IE: natural language processing (NLP), using a grammar to parse the syntax of each sentence, and statistical co-occurrence methods [1]. We focus on the latter approach, which is highly flexible and generally gives better recall, but worse precision, than NLP[18–20]. Other disadvantages of co-occurrence methods are that they are unable to extract the direction of an association and have difficulty distinguishing between direct and indirect associations [1]. However, neither of these disadvantages are important with respect to extracting disease–gene associations.

Almost all co-occurrence methods implement a frequency-based scoring scheme to account for the fact that a pair of entities or concepts may co-occur a few times without being in any way related [19,21,22]. These scoring schemes have traditionally counted either the number of sentences or the number of abstracts in which the pair co-occurred, and both sizes of text units have merit [18]. We have therefore recently introduced a scoring scheme that simultaneously takes into account both sentence-level and abstract-level co-occurrences [23].

Disease–gene associations extracted from Medline abstracts can already be searched through generalized co-occurrence tools such as CoPub [20,24] and FACTA+ [22,25]. However, as these resources are technology-centric — focusing on text mining — they do not take into account any other types of evidence. This limitation is aggravated by the fact that neither resource allows bulk download of all associations, making it difficult for others to integrate additional evidence.

### Disease–gene association databases

Several existing databases focus on or contain disease–gene associations, mainly obtained through manual curation of the biomedical literature. Unfortunately, most of these use an in-house controlled vocabulary of diseases and are subject to restrictive licenses, which makes it difficult to integrate them both from a technical and from a legal standpoint. The oldest and most famous of databases is Online Mendelian Inheritance in Man (OMIM; http://omim.org). More recent efforts include the Human Gene Mutation Database (HGMD) [26], the Comparative Toxicogenomics Database (CTD) (http://ctdbase.org/) [27,28], and Genetics Home Reference (GHR; http://ghr.nlm.nih.gov). In addition to these dedicated disease–gene association databases, UniProtKB also annotates diseases associated with each gene [9].

Databases also exist that deal with specific diseases or types of diseases, most notably cancer. The Catalog of Somatic Mutations In Cancer (COSMIC) is the most comprehensive source of information on somatic mutations and their frequencies in human cancers [29]. Mutation data is manually curated from the primary literature and annotated according to a histology and tissue ontology.

Over the last decade, genome-wide association studies (GWAS) have produced data on thousands of single nucleotide polymorphisms (SNPs) associated with the risk of hundreds of diseases. GWAS data are, however, non-trivial to work with for the non-expert, because they identify marker SNPs that are often not the actual causal SNPs [30,31]. For this reason GWAS results must be analyzed in the context of linkage disequilibrium (LD), which is defined as the non-random association of variants at two or more loci [31,32]. GWAS Central (http://www.gwascentral.org/) is a centralized database that collects the results from genetic association studies [33]. Unfortunately it provides data only for small- to medium-scale investigations and explicitly forbids using the data to create similar public resources. By contrast, the National Human Genome Research Institute (NHGRI) GWAS Catalog (http://www.genome.gov/gwastudies/) is public domain [34]. The latter is thus the basis for the derived databases DistiLD [35] and GWASdb [36] databases, which show disease-associated SNPs and genes in their chromosomal context.

Here we describe the DISEASES resource, which aims to be the most comprehensive freely available database of disease–gene associations. To this end, we have developed open-source text-mining software that performs NER of diseases and human genes as well as IE of disease–gene associations. We integrate the associations extracted through automatic text mining with evidence from databases with permissive licenses, namely manually curated associations from GHR and UniProtKB, GWAS results from DistiLD, and mutation data from COSMIC. To make the data easy to use for large-scale analyses, we map all sources of evidence to common identifiers, assign them comparable quality scores, and make them available for bulk download. We also make the information available as a user-friendly web resource (http://diseases.jensenlab.org) aimed at end users interested in individual diseases or genes.

## Material and methods

### Dictionary construction

For human gene and protein names, we used the alias file from STRING v9.1 [23], which integrates names from Ensembl [37], UniProtKB [9], and HGNC [8]. We orthographically expanded the gene symbols with the prefix ‘h’, which means *human* and is commonly used in the literature to disambiguate a human gene from its identically named orthologs in model organisms.

To construct a dictionary of diseases for use in NER, we extracted all names and synonyms from the Disease Ontology [15]. Comparing these to the dictionary of human gene names revealed that the HGNC gene symbol of a disease gene was in many cases listed in Disease Ontology as a synonym for the disease in which the gene is implicated. For example, BRCA1 and BRCA2 were listed as exact synonyms for *hereditary breast ovarian cancer*. As this would be a major source of ambiguity in the combined dictionary, we explicitly filtered out disease names that are identical to HGNC gene symbols.

To improve recall, we next automatically generated variants of the disease names. Although the terms *disease*, *disorder*, and *syndrome* have separate definitions, we found that they are used inconsistently in the literature when part of disease names; for example, *Alzheimer’s disease* is occasionally referred to as *Alzheimer’s disorder* or *Alzheimer’s syndrome*. To address this we automatically generate the two other variants if either of them is in the dictionary. Similarly, the adjectives *hereditary* and *familial* are used interchangeably, and we thus automatically replace one with the other. We also removed words in parentheses and brackets occurring at the end of disease names, unless this would cause ambiguity.

### Recognition of gene and disease names in text

To match a document against the dictionary, we have developed a highly efficient tagging algorithm, which is implemented in C++. The algorithm is described in full detail elsewhere [38], but is summarized here for completeness. Tests of the tagging speed and memory efficiency of the implementation compared to another popular tagger are also provided in our earlier publication [38].

We first tokenize the text on white space characters and special characters, such as hyphen and slash, and identify the leftmost longest matches by looking up all substrings consisting of up to 15 consecutive tokens. To make these lookups fast while handling character case variation as well as spacing and hyphenation of multiwords, we used a custom hash table to store the dictionary. The hash table is case insensitive, disregards white space characters and hyphens within name, and trims off other punctuation characters, such as quotes and parentheses, at the beginning and end of names. To match also acronyms that are not in the dictionary, we use a regular expression to search definitions of acronyms within the text and look up their long forms in the dictionary. Crucially, we globally block tagging names that would otherwise give rise to many false positives by manually inspecting the tagging results of all names that occur more than 2000 times in Medline. Many of the blocked names are acronyms; for example, the acronym for *disseminated intravascular coagulation* is DIC, which can also mean *deviance information criteria*, *differential interference contrast*, and *dissolved inorganic carbon*. By keeping track of all names that we have inspected — whether they were blocked or not — we are able to efficiently update the list of blocked names as both Medline and the dictionary grows. For each name recognized in the text we normalize it to the corresponding unique identifier and, in case of diseases, backtrack the term to the root of the ontology through is_a relationships to assign also the identifiers of all parent terms.

### Extraction and scoring of disease–gene associations

We score associations between proteins and diseases using the scoring scheme previously described [39], which is also the basis for the co-occurrence-based text-mining scores in STRING v9.1 [23] and COMPARTMENTS [40]. For completeness we reiterate the scoring scheme here.

An important feature of the scoring scheme is that it simultaneously takes into account co-occurrences at the level of abstracts as well as individual sentences. To this end, we first calculate a weighted count (*C*(*G, D*)) for each pair of a gene (*G*) and a disease (*D*) over the n abstracts in the text corpus:

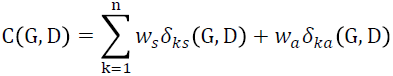

where *w_a_* = 3 and *w_s_* = 0.2 are the weights for co-occurrence within the same abstract and the same sentence, respectively, and the delta functions *δ_ak_*(*G, D*), and *δ_sk_* (*G, D*) signify whether or not *G* and *D* co-occur in abstract *k* or a sentence within it. A co-occurrence score (*S*(*G*,*D*)) is calculated from the weighted counts as:

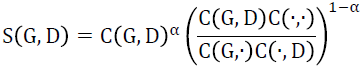

where C G, · is the sum over all diseases paired with gene *G* C(·, D) is the sum over all genes paired with disease *D*, the normalizing factor C (·,·) is the sum over all pairs of genes and diseases, and the weighting factor α = 0.6. All parameters (*w_a_*, *w_s_* and α) have in earlier work been optimized to give the best possible performance on finding functionally associated genes [23]. An important property of this function is that it not only rewards for the gene and disease being mentioned together, but also penalizes for them being frequently mentioned together with other diseases or genes, respectively.

We next convert the co-occurrence scores (*S*(*G,D*)) to z-scores (*Z*(*G,D*)), which are easier to interpret and are robust to changes in the size of the text corpus. We assume that the empirically observed score distribution is a mixture of the true signal and a lower-scoring random background, which we model as a Gaussian distribution. The full details of this score conversion have been published elsewhere [39]. Finally, we calculate the confidence score (stars) as *Z*(*G,D*)/2, limited to a maximum of four stars to account for automatic text mining never being as reliable as manually curated annotations.

### Integration of curated knowledge

The GHR database does not provide download files for use in large-scale analyses. We thus used an automated crawler to download the web page for each disease and store the disease name, which is part of the uniform resource locator (URL), along with any gene symbols listed on the web page. We were able to map the names of 390 diseases to Disease Ontology using the dictionary we developed for text mining. The pages are regularly recrawled to update with new associations; the numbers used in the manuscript are based on what was downloaded on May 31, 2013.

In case of UniProtKB, associations to diseases can be found in the KW lines through the use of 149 keywords from the UniProtKB controlled vocabulary of keywords. We were able to manually map 132 of the 149 disease keywords to corresponding concepts in the Disease Ontology. Most of the keywords that we could not map, such as *Disease mutation*, were not disease names.

We mapped HGNC gene symbols from GHR and identifiers from UniProtKB to their identifiers in STRING v9.1 using the alias file [23]. We subsequently used the explicitly annotated disease–gene associations from GHR and UniProtKB to infer broader Disease Ontology concepts via the is_a relationships in the ontology. As all disease– gene annotations imported and inferred from the two databases are based on manual curation, we assigned them a confidence score of five stars.

### Benchmark of text-mining results

To assess the quality of the text-mining results, we constructed a reference set based on the manually curated annotations imported from GHR and UniProtKB. Due to the hierarchical nature of the Disease Ontology, it is necessary to select on a subset of terms to be used as the basis for the assessment. To this end, we chose to use the subset of terms that were explicitly annotated in the two databases (as opposed to inferred through is_a relationships). In case one term was a child term of another, we selected the broader parent term. This resulted in a positive reference set of 2780 associations between 2001 genes and 173 diseases. We defined the negative set as all other 343393 possible pairings of the same genes and diseases.

We next sorted the text-mined associations descending by score and compared them to the reference set. We present the results as receiver operating characteristic (ROC) curves by plotting the true positive rate (TPR) as function of false positive rate (FPR), considering either all disease–gene associations or only the best-scoring association per gene (Figure 1). We compare these results to two random backgrounds. One is simple random shuffling of the disease–gene pairs, which ignores that some diseases are associated with many more genes than others. To correct for this, the second random background is calculated by sorting the disease–gene pairs descending by prior probability of the disease. Because the prior of each disease is estimated based on the reference set itself, this likely overestimates the performance that can be attained by random guessing.

**Figure 1:**
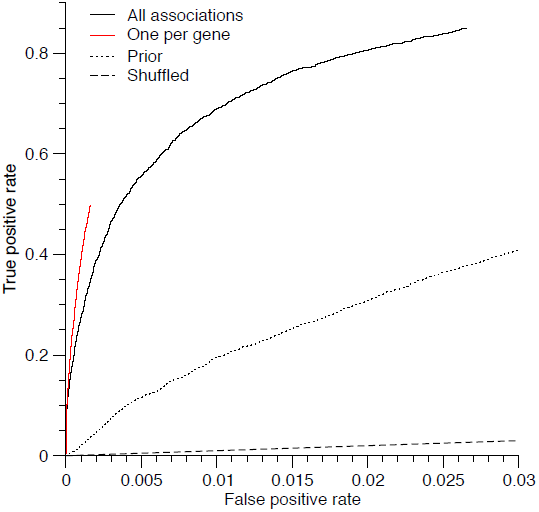
Benchmark of disease–gene associations obtained through text mining. The receiver operating characteristic (ROC) curves shows the true positive rate (TPR) as function of false positive rate (FPR) when considering all associations (black) and when considering only the highest scoring association for each gene (red). The dashed and dotted curves show the random expectations according to simple shuffling and prior-based ranking, respectively. The curves do not intercept TPR = 1 and FPR = 1, because some disease–gene pairs in the benchmark set are not found mentioned together in Medline, for which reason they have no text-mining score.

### Integration of mutation and GWAS data

To integrate cancer mutation data from COSMIC [29], we manually created mappings between terms listed in the fields “Site primary” and “Histology” and Disease Ontology concepts classified under “organ system cancer” and “cell type cancer”, respectively. We mapped the genes to STRING v9.1 identifiers via the Ensembl transcript identifiers provided by COSMIC. For each pair of a gene (G) and a disease (D) we counted the number of disease samples carrying at least one somatic missense or nonsense mutation within the gene (N(G, D)). We discarded pairs with a count less than 10 and derived confidence scores (stars) as log_10_ (N(G, D)) − 0.5, limiting it to at most four stars.

To include also GWAS data, we integrated information from the DistiLD database [35], which maps genes and disease-associated SNPs onto so-called LD blocks defined based on data from the HapMap Project [41]. We assigned each SNP with a p-value less than 10^−5^ to the nearest gene within the same LD block. The “Disease/Trait” descriptors from the NHGRI GWAS Catalog were mapped to the corresponding Disease Ontology concepts through the ICD-10 annotations from DistiLD, the Disease Ontology Lite annotations from GWASdb [36], and manual inspection of conflicts. The resulting disease–gene associations were assigned a confidence score (stars) using the formula 3 – log_10_(max(P, P_min_)) where P is the p-value, P_min_ is the genome-wide GWAS significance threshold (5 ⋅ 10^−8^).

## Results and discussion

### Dictionary-based tagger software

We have developed a highly efficient NER method for diseases and human genes, which are normalized to identifiers from Disease Ontology [15] and STRING v9.1 [23], respectively. On a server with two Intel E5520 processors and 24GB of random access memory (RAM), starting the tagger and loading the dictionary took only 4.2 seconds. Once started, the tagger used 260MB of RAM and was able to process 360 Medline abstracts per second on a single processor core (measured on a corpus of 100,000 Medline abstracts). The tagger software bundled with a dictionary of disease and human gene names is available for download under the BSD license.

### Cooccurrence-based disease–gene associations

Because the NER task is for us only a step on the way towards the goal of extracting disease–gene associations, we chose to focus our benchmarking effort on assessing the quality of the end result. We therefore compared the text-mined associations to the manually curated associations imported from GHR and UniProtKB in two ways: 1) considering all disease–gene associations, and 2) considering only the highest scoring disease for each gene. The results of these comparisons (Figure 1) show that our text-mining system is able to extract a large fraction of the known disease–gene associations with high specificity (low FPR). If a user were to simply trust the highest scoring disease association for each gene, 50% of all manually curated disease–gene associations in the benchmark set would be found at a FPR of only 0.16%.

The high quality of text-mining results is reflected by the fact that they are already being used extensively. The text-mined associations from DISEASES are included in the widely used GeneCards database [42]. They have also been used as a basis for inference of disease associations for miRNAs from their predicted target genes [39] and for enrichment analysis of autism-related genes [43].

### Contents of the database

Although we have in this paper placed most emphasis on the text-mining aspects, the DISEASES database integrates disease–gene associations from several sources. This is advantageous, because every source of associations has its shortcomings. Table 1 provides an overview of the total evidence landscape of the database, showing that the text-mining pipeline is indeed the largest single contributor of associations. However, it is important to note that this number depends strongly on the confidence cutoff; indeed the number of associations obtained from the manually curated databases rivals the number of text-mined associations with at least 3 confidence stars. Mutation data from COSMIC and GWAS data from DistiLD also both contribute a sizeable number of associations; however, the former data source only relates genes to cancers.

**Table 1:** **Overview of disease–gene association evidence.** Each row shows the number of genes, diseases and associations between them that are supported by a given type, confidence level (in case of Text mining), or source (in case of Knowledge and Experiments). The numbers in parentheses specify the counts prior to backtracking of Disease Ontology terms through is_a relationships.

| **Evidence** | **Genes** | **Diseases** | **Associations** |
| --- | --- | --- | --- |
| Text mining | 15631 | 4598 | 478407 |
| - 4 star confidence | 478 | 662 | 1044 |
| - 3 star confidence | 3207 | 2267 | 15226 |
| - 2 star confidence | 12706 | 4354 | 142892 |
| Knowledge | 2001 | 735<br>(453) | 15231<br>(2953) |
| - Genetics Home Reference | 965 | 671<br>(390) | 7551<br>(1169) |
| - UniProtKB | 1651 | 271<br>(120) | 11576<br>(2187) |
| Experiments | 10711 | 423<br>(264) | 89073<br>(20206) |
| - COSMIC | 8786 | 142<br>(76) | 55791<br>(13050) |
| - DistiLD | 4315 | 351<br>(210) | 36650<br>(7185) |
| Total | 17606 | 4610 | 543405 |

All disease–gene associations from all evidence sources are available for bulk download in tab-delimited format under the Creative Commons Attribution (CC-BY) license.

### The DISEASES web interface

Whereas tab-delimited files are convenient for bioinformaticians wanting to perform large-scale analyses or create derived resources, a user-friendly web interface better caters to researchers interested in individual genes or diseases.

We have thus developed a web interface for the DISEASES resource that allows users to either query for a gene to find associated diseases or query for a disease to find associated genes (Figure 2). In either case, the user will be presented with three tables called Knowledge, Experiments, and Text mining. These show the manually curated associations from GHR and UniProtKB, the mutation and association data from COSMIC and DistiLD, and the text-mined associations, respectively. Besides summarizing the imported information, the Knowledge and Experiments tables provide direct hyperlinks to the source entries in the external databases.

**Figure 2:**
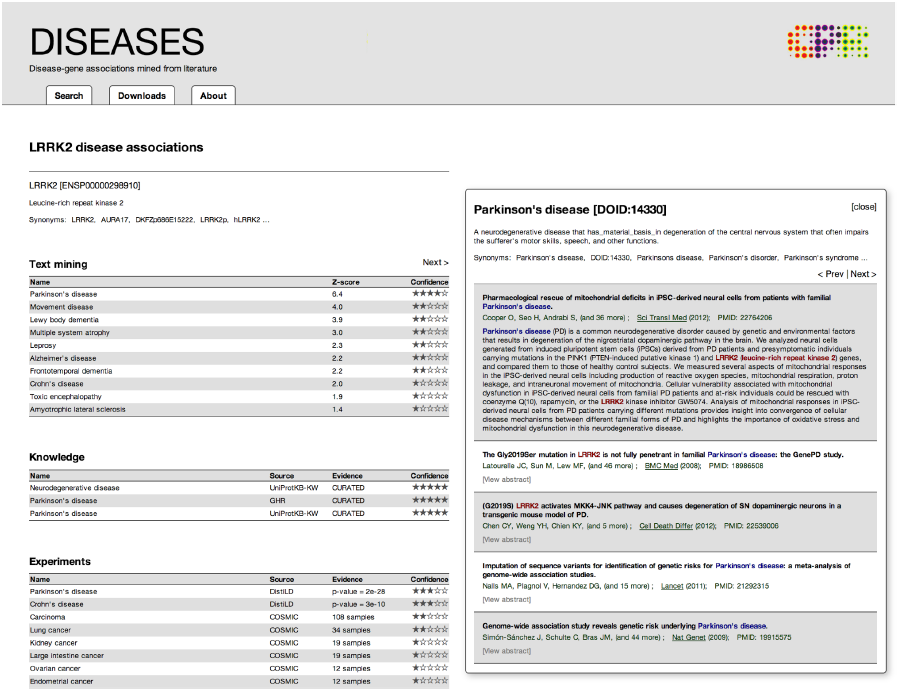
The DISEASES web resource. The figure shows how the disease–gene associations are presented in the web interface, exemplified by the LRRK2 gene. The three tables provide the user with an overview of the evidence from text mining, curated knowledge, and experimental data. Clicking on an association, e.g. to Parkinson’s disease, in the Text mining table gives access to the underlying abstracts with the co-occurring gene and disease highlighted. The two other tables provide hyperlinks to the relevant entries in the source databases.

The table summarizing the text-mined evidence deserves special attention. As the text-mining method correctly takes into account information from the narrower child terms of each disease, the text-mined disease associations for a gene have inherent redundancy. When showing the list of diseases associated with a gene of interest, the web interface thus dynamically filters out redundant Disease Ontology terms for which better alternatives are present. The web interface also gives the user the possibility to inspect the text-mining evidence behind any disease–gene association by viewing the underlying abstracts with the gene and disease names highlighted.

### Generality of the approach

The approach to text mining described in this paper is readily applicable to recognize other types of named entities in text and extract associations among them. Using the same tagger with a dictionary constructed from the NCBI Taxonomy [44], we were able to accurately identify taxonomic names in the biomedical literature [38]. We are currently extending that work to identify environments from the Environment Ontology [45] in text, for example, from the Encyclopedia of Life [46]. We have even used a slightly modified version of the tagger as part of a method for recognition of adverse drug events in Danish clinical narratives [47]. This illustrates the flexibility of a simple dictionary-based NER approach in terms of applicability to new knowledge domains.

Combining the tagger with the co-occurrence scoring scheme for the purpose of IE is equally flexible. As previously mentioned, the scoring scheme was originally developed to extract functional associations between proteins for use in the STRING database based on co-occurrence of gene names within biomedical literature [23]. In addition to using it for disease–gene associations as described here, we have since applied the same scoring scheme to extract information on protein–small molecule associations in the STITCH database [48], protein subcellular localization in the COMPARTMENTS database [40], and tissue distribution of proteins in the TISSUES database (http://tissues.jensenlab.org).

Besides using the same methods for NER and IE, DISEASES and the other resources mentioned above have in common that they integrate heterogeneous evidence from many sources. This sets them aside from the many resources that use text mining to extract associations between a wide variety of named entities and concepts. As tool developers, it is easiest and most efficient to be technology-centric and apply a single technology, such as text mining, to a wide range of topics. However, from a user’s perspective, a resource that integrates many sources of information pertaining to a single topic of interest is usually what is sought after. We attempt to find a compromise by creating a general framework, which allows us to set up resources that each integrate information on a different topic but are maintainable, because they share software infrastructure.

## Conclusions

We have developed a dictionary-based NER tool for Disease Ontology concepts and combined it with a co-occurrence scoring scheme to efficiently and accurately extract disease–gene associations from Medline. We have integrated these with manually curated associations from the GHR and UniProtKB databases as well as somatic mutation and GWAS data from COSMIC and DistiLD, respectively. We make the resulting database available as a searchable user-friendly web resource at http://diseases.jensenlab.org, where bulk datasets and the NER software are also available for download.

## Acknowledgments

This work was in part funded by the Novo Nordisk Foundation Center for Protein Research and by the European Union’s Seventh Framework Programme (FP7/2007-2013) under grant agreement n259348. The authors thank Andreas Bok Andersen for help with developing the web interface.

## References

[1] L.J. Jensen, J. Saric, P. Bork, Literature mining for the biologist: from information retrieval to biological discovery., Nat. Rev. Genet. 7 (2006) 119–29. doi:10.1038/nrg1768.

[2] L. Chen, H. Liu, C. Friedman, Gene name ambiguity of eukaryotic nomenclatures., Bioinformatics. 21 (2005) 248–56. doi:10.1093/bioinformatics/bth496.

[3] K. Fukuda, A. Tamura, T. Tsunoda, T. Takagi, Toward information extraction: identifying protein names from biological papers., Pac. Symp. Biocomput. (1998) 707–18. http://www.ncbi.nlm.nih.gov/pubmed/9697224 (accessed January 16, 2014).

[4] B. Settles, ABNER: an open source tool for automatically tagging genes, proteins and other entity names in text., Bioinformatics. 21 (2005) 3191–2. doi:10.1093/bioinformatics/bti475.

[5] G. Zhou, D. Shen, J. Zhang, J. Su, S. Tan, Recognition of protein/gene names from text using an ensemble of classifiers., BMC Bioinformatics. 6 Suppl 1 (2005) S7. doi:10.1186/1471-2105-6-S1-S7.

[6] D. Hanisch, K. Fundel, H.-T. Mevissen, R. Zimmer, J. Fluck, ProMiner: rule-based protein and gene entity recognition., BMC Bioinformatics. 6 Suppl 1 (2005) S14. doi:10.1186/1471-2105-6-S1-S14.

[7] S. Gaudan, H. Kirsch, D. Rebholz-Schuhmann, Resolving abbreviations to their senses in Medline., Bioinformatics. 21 (2005) 3658–64. doi:10.1093/bioinformatics/bti586.

[8] K.A. Gray, L.C. Daugherty, S.M. Gordon, R.L. Seal, M.W. Wright, E.A. Bruford, Genenames.org: the HGNC resources in 2013., Nucleic Acids Res. 41 (2013) D545–52. doi:10.1093/nar/gks1066.

[9] The UniProt Consortium, Activities at the Universal Protein Resource (UniProt)., Nucleic Acids Res. 42 (2014) D191–8. doi:10.1093/nar/gkt1140.

[10] P.B. Jensen, L.J. Jensen, S. Brunak, Mining electronic health records: towards better research applications and clinical care., Nat. Rev. Genet. 13 (2012) 395–405. doi:10.1038/nrg3208.

[11] F.S. Roque, P.B. Jensen, H. Schmock, M. Dalgaard, M. Andreatta, T. Hansen, et al., Using electronic patient records to discover disease correlations and stratify patient cohorts., PLoS Comput. Biol. 7 (2011) e1002141. doi:10.1371/journal.pcbi.1002141.

[12] C. Friedman, H. Liu, L. Shagina, S. Johnson, G. Hripcsak, Evaluating the UMLS as a source of lexical knowledge for medical language processing., Proc. AMIA Symp. (2001) 189–93. http://www.pubmedcentral.nih.gov/articlerender.fcgi?artid=2243298&tool=pmcentrez&rendertype=abstract.

[13] G.K. Savova, J.J. Masanz, P. V Ogren, J. Zheng, S. Sohn, K.C. Kipper-Schuler, et al., Mayo clinical Text Analysis and Knowledge Extraction System (cTAKES): architecture, component evaluation and applications., J. Am. Med. Inform. Assoc. 17 (2010) 507–13. doi:10.1136/jamia.2009.001560.

[14] H. Kilicoglu, G. Rosemblat, M. Fiszman, T.C. Rindflesch, Constructing a semantic predication gold standard from the biomedical literature., BMC Bioinformatics. 12 (2011) 486. doi:10.1186/1471-2105-12-486.

[15] L.M. Schriml, C. Arze, S. Nadendla, Y.-W.W. Chang, M. Mazaitis, V. Felix, et al., Disease Ontology: a backbone for disease semantic integration., Nucleic Acids Res. 40 (2012) D940–6. doi:10.1093/nar/gkr972.

[16] B. Smith, M. Ashburner, C. Rosse, J. Bard, W. Bug, W. Ceusters, et al., The OBO Foundry: coordinated evolution of ontologies to support biomedical data integration., Nat. Biotechnol. 25 (2007) 1251–5. doi:10.1038/nbt1346.

[17] J.D. Osborne, J. Flatow, M. Holko, S.M. Lin, W.A. Kibbe, L.J. Zhu, et al., Annotating the human genome with Disease Ontology., BMC Genomics. 10 Suppl 1 (2009) S6. doi:10.1186/1471-2164-10-S1-S6.

[18] J. Ding, D. Berleant, D. Nettleton, E. Wurtele, Mining MEDLINE: abstracts, sentences, or phrases?, Pac. Symp. Biocomput. (2002) 326–37. http://www.ncbi.nlm.nih.gov/pubmed/11928487 (accessed January 16, 2014).

[19] J.D. Wren, H.R. Garner, Shared relationship analysis: ranking set cohesion and commonalities within a literature-derived relationship network, Bioinformatics. 20 (2004) 191–198. doi:10.1093/bioinformatics/btg390.

[20] B.T.F. Alako, A. Veldhoven, S. van Baal, R. Jelier, S. Verhoeven, T. Rullmann, et al., CoPub Mapper: mining MEDLINE based on search term co-publication., BMC Bioinformatics. 6 (2005) 51. doi:10.1186/1471-2105-6-51.

[21] T.K. Jenssen, a Laegreid, J. Komorowski, E. Hovig, A literature network of human genes for high-throughput analysis of gene expression., Nat. Genet. 28 (2001) 21–8. doi:10.1038/88213.

[22] Y. Tsuruoka, J. Tsujii, S. Ananiadou, FACTA: a text search engine for finding associated biomedical concepts., Bioinformatics. 24 (2008) 2559–60. doi:10.1093/bioinformatics/btn469.

[23] A. Franceschini, D. Szklarczyk, S. Frankild, M. Kuhn, M. Simonovic, A. Roth, et al., STRING v9.1: protein-protein interaction networks, with increased coverage and integration., Nucleic Acids Res. 41 (2013) D808– 15. doi:10.1093/nar/gks1094.

[24] W.W.M. Fleuren, S. Verhoeven, R. Frijters, B. Heupers, J. Polman, R. van Schaik, et al., CoPub update: CoPub 5.0 a text mining system to answer biological questions., Nucleic Acids Res. 39 (2011) W450–4. doi:10.1093/nar/gkr310.

[25] Y. Tsuruoka, M. Miwa, K. Hamamoto, J. Tsujii, S. Ananiadou, Discovering and visualizing indirect associations between biomedical concepts., Bioinformatics. 27 (2011) i111–9. doi:10.1093/bioinformatics/btr214.

[26] P.D. Stenson, M. Mort, E. V Ball, K. Shaw, A.D. Phillips, D.N. Cooper, The Human Gene Mutation Database: building a comprehensive mutation repository for clinical and molecular genetics, diagnostic testing and personalized genomic medicine., Hum. Genet. (2013). doi:10.1007/s00439-013-1358-4.

[27] A.P. Davis, C.G. Murphy, R. Johnson, J.M. Lay, K. Lennon-Hopkins, C. Saraceni-Richards, et al., The Comparative Toxicogenomics Database: update 2013., Nucleic Acids Res. 41 (2013) D1104–14. doi:10.1093/nar/gks994.

[28] A.P. Davis, T.C. Wiegers, P.M. Roberts, B.L. King, J.M. Lay, K. Lennon-Hopkins, et al., A CTD-Pfizer collaboration: manual curation of 88,000 scientific articles text mined for drug-disease and drug-phenotype interactions., Database (Oxford). 2013 (2013) bat080. doi:10.1093/database/bat080.

[29] S.A. Forbes, G. Bhamra, S. Bamford, E. Dawson, C. Kok, J. Clements, et al., The Catalogue of Somatic Mutations in Cancer (COSMIC)., Curr. Protoc. Hum. Genet. Chapter 10 (2008) Unit 10.11. doi:10.1002/0471142905.hg1011s57.

[30] M.I. McCarthy, G.R. Abecasis, L.R. Cardon, D.B. Goldstein, J. Little, J.P.A. Ioannidis, et al., Genome-wide association studies for complex traits: consensus, uncertainty and challenges., Nat. Rev. Genet. 9 (2008) 356–69. doi:10.1038/nrg2344.

[31] M. Slatkin, Linkage disequilibrium--understanding the evolutionary past and mapping the medical future., Nat. Rev. Genet. 9 (2008) 477–85. doi:10.1038/nrg2361.

[32] D. Altshuler, M.J. Daly, E.S. Lander, Genetic mapping in human disease., Science. 322 (2008) 881–8. doi:10.1126/science.1156409.

[33] G.A. Thorisson, O. Lancaster, R.C. Free, R.K. Hastings, P. Sarmah, D. Dash, et al., HGVbaseG2P: a central genetic association database., Nucleic Acids Res. 37 (2009) D797–802. doi:10.1093/nar/gkn748.

[34] L.A. Hindorff, P. Sethupathy, H.A. Junkins, E.M. Ramos, J.P. Mehta, F.S. Collins, et al., Potential etiologic and functional implications of genome-wide association loci for human diseases and traits., Proc. Natl. Acad. Sci. U. S. A. 106 (2009) 9362–7. doi:10.1073/pnas.0903103106.

[35] A. Pallejà, H. Horn, S. Eliasson, L.J. Jensen, DistiLD Database: diseases and traits in linkage disequilibrium blocks., Nucleic Acids Res. 40 (2012) D1036–40. doi:10.1093/nar/gkr899.

[36] M.J. Li, P. Wang, X. Liu, E.L. Lim, Z. Wang, M. Yeager, et al., GWASdb: a database for human genetic variants identified by genome-wide association studies., Nucleic Acids Res. 40 (2012) D1047–54. doi:10.1093/nar/gkr1182.

[37] P. Flicek, I. Ahmed, M.R. Amode, D. Barrell, K. Beal, S. Brent, et al., Ensembl 2013., Nucleic Acids Res. 41 (2013) D48–55. doi:10.1093/nar/gks1236.

[38] E. Pafilis, S.P. Frankild, L. Fanini, S. Faulwetter, C. Pavloudi, A. Vasileiadou, et al., The SPECIES and ORGANISMS Resources for Fast and Accurate Identification of Taxonomic Names in Text., PLoS One. 8 (2013) e65390. doi:10.1371/journal.pone.0065390.

[39] S. Mørk, S. Pletscher-Frankild, A. Palleja Caro, J. Gorodkin, L.J. Jensen, Protein-driven inference of miRNA-disease associations., Bioinformatics. (2013). doi:10.1093/bioinformatics/btt677.

[40] J.X. Binder, S. Pletscher-Frankild, K. Tsafou, C. Stolte, S.I. O’Donoghue, R. Schneider, et al., COMPARTMENTS: unification and visualization of protein subcellular localization evidence., Database (Oxford). 2014 (2014) bau012. doi:10.1093/database/bau012.

[41] The International HapMap Consortium, A haplotype map of the human genome., Nature. 437 (2005) 1299–320. doi:10.1038/nature04226.

[42] M. Safran, I. Dalah, J. Alexander, N. Rosen, T. Iny Stein, M. Shmoish, et al., GeneCards Version 3: the human gene integrator., Database (Oxford). 2010 (2010) baq020. doi:10.1093/database/baq020.

[43] B.E. Eisinger, M.C. Saul, T.M. Driessen, S.C. Gammie, Development of a versatile enrichment analysis tool reveals associations between the maternal brain and mental health disorders, including autism., BMC Neurosci. 14 (2013) 147. doi:10.1186/1471-2202-14-147.

[44] E.W. Sayers, T. Barrett, D.A. Benson, S.H. Bryant, K. Canese, V. Chetvernin, et al., Database resources of the National Center for Biotechnology Information., Nucleic Acids Res. 37 (2009) D5–15. doi:10.1093/nar/gkn741.

[45] P.L. Buttigieg, N. Morrison, B. Smith, C.J. Mungall, S.E. Lewis, The environment ontology: contextualising biological and biomedical entities., J. Biomed. Semantics. 4 (2013) 43. doi:10.1186/2041-1480-4-43.

[46] E.O. Wilson, The encyclopedia of life, Trends Ecol. Evol. 18 (2003) 77–80. doi:10.1016/S0169-5347(02)00040-X.

[47] R. Eriksson, P.B. Jensen, S. Frankild, L.J. Jensen, S. Brunak, Dictionary construction and identification of possible adverse drug events in Danish clinical narrative text., J. Am. Med. Inform. Assoc. 20 (2013) 947–53. doi:10.1136/amiajnl-2013-001708.

[48] M. Kuhn, D. Szklarczyk, S. Pletscher-Frankild, T.H. Blicher, C. von Mering, L.J. Jensen, et al., STITCH 4: integration of protein-chemical interactions with user data., Nucleic Acids Res. 42 (2014) D401–7. doi:10.1093/nar/gkt1207.

